# Gene network topology drives the mutational landscape of gene expression

**DOI:** 10.1101/2024.11.28.625874

**Authors:** Sylvain Pouzet, Arnaud Le Rouzic

**Affiliations:** Université Paris-Saclay, CNRS, IRD, UMR Evolution, Génomes, Comportement et Ecologie, Gif-Sur-Yvette, France; Institut Curie, Université PSL, Sorbonne Université, CNRS UMR168, Laboratoire Physico Chimie Curie, Paris, France

**Keywords:** Gene regulatory networks, Distribution of fitness effects, Population genetics model, Evolutionary systems biology

## Abstract

Regulatory mutations, coding sequence alterations, and gene deletions and duplications are generally expected to have qualitatively different effects on fitness. We aim to ground this expectation within a theoretical framework using evolutionary simulations of gene regulatory networks (GRNs) controlling the expression of fitness-related genes. We examined the distribution of fitness effects as a function of the type of mutation and the topology of the gene network. Contrary to our expectation, the GRN topology had more influence on the effect of mutations than the type of mutation itself. In particular, the topology conditioned (i) the speed of adaptation, (ii) the distribution of fitness effects, and (iii) the degree of pleiotropy, acting as explanatory factor for all mutation types. All mutations had the potential to participate in adaptation, although their propensity to generate beneficial variants differed according to the network topology. In scalefree networks, the most common topology for biological networks, coding mutations were more pleiotropic and overrepresented in both beneficial and deleterious mutations, while regulatory mutations were more often neutral. However, this observation was not general, as this pattern was reversed in alternative networks. These results highlight the critical role of network topology in defining mutations’ contributions to adaptation.

---

The nature of the molecular variations that feed adaptive evolution is a long-standing question in evolutionary biology (King and Wilson 1975; Stern and Orgogozo 2008). Mutations recruited in the course of evolution may affect DNA in different ways: some mutations change gene coding sequences, and thus the functional features of the gene products, others affect gene expression, *i.e.* essentially, the amount of RNA and proteins produced by the genes. Likewise, gene duplications or deletions change the structure and complexity of the biological systems and, ultimately, the number of dimensions in which genetic evolution takes place. There is compelling evidence that all kinds of mutations are utterly important for adaptive evolution, but the circumstances in which different mutations are more likely to confer a fitness advantage remain largely elusive.

Morphological and physiological variation generally roots into large and complex regulatory networks that control the pattern of gene expression in space and time (Carroll 2008; Stern and Orgogozo 2008). As most complex genetic systems, gene regulatory networks (GRNs) evolve through regulatory or coding mutations, as well as gene duplications and deletions. This mutational inflow, filtered out by negative (conservative) and positive (adaptive) selection, conditions the GRN structure and evolvability (Aguilar-Rodríguez and Payne 2021). Gene expression can be affected by mutations altering or disrupting regulatory sites in the promoters of genes (Moses et al. 2006; Prud’homme et al. 2007), or creating new binding sites (Yona et al. 2018). Changes in gene expression drive adaptation in a wide variety of species (Fraser 2013; López-Maury et al. 2008). Adaptive mutations in the exons of protein-coding genes have been for a long time the only available data on molecular evolution (Sawyer et al. 2003), and countless examples of adaptive coding sequence mutations are known, affecting morphological traits but also gene expression (Hoekstra and Coyne 2007; G. P. Wagner and V. J. Lynch 2008). Indeed, non-synonymous mutations in the coding region of Transcription Factors (TFs) can change their efficiency, binding affinity, or target DNA sequence (Galant and Carroll 2002; M. Lynch 2007; Ronshaugen et al. 2002). Finally, the role of gene duplications in adaptation and evolutionary innovation has also been convincingly documented (Kondrashov 2012; Qian and J. Zhang 2014), and the long-term evolution of GRNs is thought to occur through TF duplication and deletion events, which also drives the evolution of the network size and shape (Teichmann and Babu 2004; J. Zhang 2003).

Yet, beyond the fact that any kind of mutation could at least occasionally fuel adaptive change, their effects on fitness are not necessarily the same: duplications and deletions may have large-scale effects, mutations in the coding sequence of TFs may also affect in *trans* a substantial amount of target genes, while *cis*-acting mutations tend to produce more localized effects. The consequences of mutations on the expression and the coding sequence of transcription factors probably also depend on structure of gene networks (Stern and Orgogozo 2009; Vande Zande and Wittkopp 2022), e.g., the number of downstream connected genes, or the presence of feedback or feedforward loops that can amplify or buffer the effects of the mutations.

The network rewiring, involving the evolution of promoter regions, and the ability for TFs to bind and regulate gene expression, is often considered the primary mode of GRN evolution (Carroll 2008; Wittkopp and Kalay 2012). The ”*cis*-regulatory hypothesis” compellingly argues that regulatory mutations in gene promoters are the most frequent ones involved in morphological adaptation (Carroll 2005; Wray 2007). While this *cis*-regulatory hypothesis accumulates convincing empirical support in a wide variety of organisms and traits (Coolon et al. 2014; Krishnan et al. 2022; Marand et al. 2023), it remains a verbal model, not firmly grounded into the formal frameworks of population and quantitative genetics which require explicit Genotype-Phenotype or Genotype-Fitness maps (Pavlicev et al. 2022). Overall, why and how differences in the nature of mutations lead to variation in gene expression, phenotypic traits, and fitness remain poorly understood (Signor and Nuzhdin 2018). The vast majority of formal GRN evolution models only consider regulatory mutations (Ciliberti et al. 2007; Kaneko and Kikuchi 2022; A. Wagner 1996). Gene duplications in GRNs have sometimes been implemented in evolutionary models, mostly to study network size and topology (Crombach and Hogeweg 2008; Kuo et al. 2006; Solé and Valverde 2008; A. Wagner 1994), and not to be compared with regulatory mutations. Yet, to the best of our knowledge, there is no formal attempt yet at comparing theoretically the genotype-to-phenotype map and the distribution of fitness effects of all major mutational mechanisms together.

We aim to address this problem by simulating the evolution of gene regulatory networks, incorporating coding mutations besides regulatory mutations and duplication/deletions, within a population genetics framework. For this, we modified Wagner’s popular GRN evolution model (inspired from (A. Wagner 1996)), so that the influence of a TF is determined by both its interaction’s strength with the promoters of other genes, and by its intrinsic activity. We allowed the networks to evolve under stabilizing selection to match a predetermined ”optimal” gene expression pattern. These networks acquired three distinct topologies (highly-connected, random and scale-free) depending on the initial conditions. Then, we simulated an environmentally-driven shift in the fitness function. This enabled us to analyze the propensity of regulatory mutations, coding mutations, gene duplications and deletions to induce deleterious, neutral, or beneficial fitness changes in the new environment for the different network topologies.

## Models and Methods

### Gene network model

The model is inspired from the framework proposed by A. Wagner (A. Wagner 1994; A. Wagner 1996), a theoretical setting in evolutionary biology (Fierst and Phillips 2015). The model consists in coupling a simplified gene network with a traditional population genetics individual-based simulation. The gene network model takes the individual genotypes as an input, and provides a vector of gene expressions as an output.

The genotype for the *n* genes of the network consisted in a *n* × *n* matrix **W**, encoding the pairwise gene interactions, and a *n*-vector **C** representing gene activities. The total regulation effect of gene *j* on gene *i* depends on *P_j_* (the concentration of *j*, scaled between 0 and 1), *C_j_*(the activity of *j*, *C_j_* ≤ 0), and *W_ij_* (the sensitivity of *i* to the transcription factor *j*), and was calculated as the product *W_ij_* × *C_j_* × *P_j_*. The elements *W_ij_* of **W** were real numbers, representing the effect of gene *j* on the expression of gene *i*; gene *j* was an inhibitor of *i* when *W_ij_ <* 0, and an activator when *W_ij_ >* 0. Each gene was a potential transcription factor (all genes could evolve to regulate one or several other genes of the network), but self-regulation was not allowed (*W_i_i* = 0) to facilitate the evolution of connected networks. The network was updated for *T* = 20 discrete time steps according to the following process (expressed with linear algebra notation):

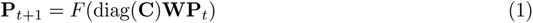

where *F* (*f* (*x*_1_), …, *f* (*x_n_*)) is a multidimensional sigmoid function,

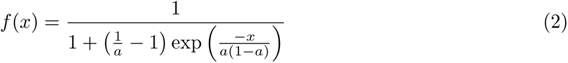

scaling its argument *x* between 0 and 1 (Fig. 1). diag(**C**) is a diagonal matrix containing the elements of the vector **C**.

**Figure 1:**
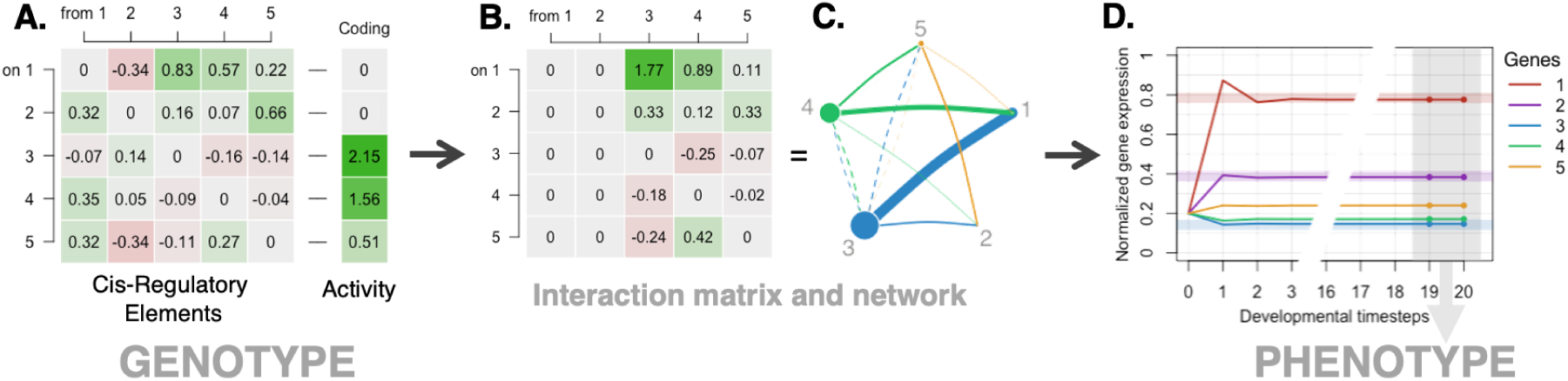
Overview of the gene network model. For clarity, the illustrated network involved 5 genes (vs. 10 in our simulations). **A.** The genotype is encoded as a 5 × 5 matrix *W* representing the regulations among genes, and an activity vector *C*. **B.** The network update operation from equation (1) is equivalent to scaling the regulatory elements interaction matrix column-wise by the coding (activity) vector, such that a gene with activity=0 results in a column set to 0 since no regulation takes place (the transcription factor is expressed but has no regulatory effect, as in e.g., genes 1 and 2 here). **C.** The interaction matrix can be represented as a graph, here showing activation (solid lines) and repression (dashed lines) of different intensities (line thickness) between genes of different activities (node sizes). **D.** From this new interaction matrix, gene expression of each gene is computed for each of the 20 discrete steps of the dynamic process (starting with constitutive expression *a* = 0.2). Final gene expression was computed by averaging the last two timesteps, and fitness was determined based on the optimum of selected genes (3 genes our of 5 shown here, thick horizontal lines).

The initial gene expression values **P**_*t*=0_ = (*a, …, a*) were set to the constitutive expression *a* = 0.2. The individual’s expression phenotype 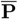 was calculated as the mean expression of each gene for the last 2 time steps of the network dynamics 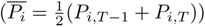.

### Population genetics

Populations consisted of *N* = 1, 000 diploid, hermaphrodite individuals, characterized by their genotype **W**, **C** and their phenotype 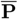. Each of the *n* loci in the genome was encoded by *n* + 1 elements (the *n* regulatory elements corresponding to a single row of **W**, and the activity *C_i_*). The genotype at heterozygote loci was obtained by averaging the values of both alleles. Loci were recombining freely; sexual reproduction consisted in merging two parental gametes, and gametes were produced by drawing an allele randomly at each locus.

Generations were non-overlapping. A generation consisted in (i) computing the phenotype 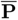 and the fitness *w* for all *N* individuals, (ii) drawing 2*N* parents with replacement from the population, with a probability proportional to their fitness, and (iii) merging these gametes into *N* new individuals for the next generation. Mutations were drawn as gametes were formed.

Fitness was calculated as *w* = *w_S_* ×*w_I_*, where *w_S_* stands for the stabilizing selection component, and *w_I_* for a network stability component. Stabilizing selection consisted in applying a bell-shaped multivariate fitness function on the expression of a subset of *n_s_* = 5 selected genes:

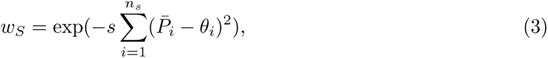

where 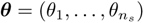 is a vector of phenotypic optima. In this setting, fitnesses were relative to the perfect phenotype (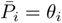 for all selected genes leads to a fitness *w_S_* = 1). The environmental change was simulated by shifting two expression optima by a deviation 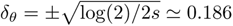, so that the fitness drops by 50% 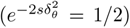 - see also Supplementary Figure S1. Stability selection consisted in applying a penalty to unstable networks which had not reached an equilibrium after *T* time steps. In practice,

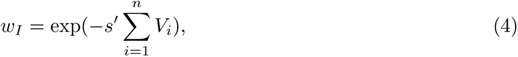

where *s*′ = 46, 000 quantifies the (strong) cost of instability (a standard deviation of 0.01 in equilibrium gene expression leads to a 99% fitness drop), 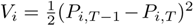 being the variance of gene expression *P_it_*computed over the two last time steps.

### Mutations

Three types of mutations were considered: regulatory mutations, coding mutations, and gene deletions or duplications. Regulatory mutation consisted in modifying a random element of the gene promoter by a deviation *δ_r_*, so that the new genotype becomes 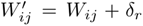. Coding mutations followed the same principle, a deviation *δ_c_* was applied to the gene activity 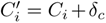. Gene activities could not be negative, and 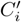 was set to 0 if the previous operation resulted in a negative activity. Gene deletion was simulated by forcing the corresponding gene expression *P_i_* to 0 at every step of the gene network dynamics. Gene duplication was simulated by duplicating a random row and its corresponding column in the matrix **W**, so that both duplicated genes were regulated by the same transcription factors, and had the same effect on other genes. The corresponding element in vector **C** was also duplicated.

Mutations were carried out in two contexts: during adaptation in the individual-based simulations, and for analysis after the simulations. Only regulatory and coding mutations were considered during adaptation. In simulations, mutations occurred with a rate *µ_r_* = 0.01 and *µ_c_* = 0.01 per gamete for regulatory and coding mutations, respectively. Mutational effects *δ_r_* and *δ_c_* were drawn from Gaussian distributions with mean 0 and variance 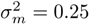. For the analysis stage, after the simulations, mutations were applied to the mean genotype of the population 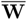 and 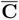, obtained by averaging the values of all 2*N* alleles at generation 1, 000. The mutational effects *δ_r_* and *δ_c_* were forced to specific values, and the number of deleted or duplicated genes was also controlled. Deleted genes were never picked among genes under selection.

### Fitness Effects

In order to determine whether a mutation is deleterious, neutral, or beneficial immediately after the environmental change, we computed a differential fitness 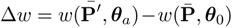, where the mutant genotype (of phenotype **P**^′^) differs from the ancestral genotype by one mutation: regulatory mutation when W was altered, coding mutation when C was altered, or duplication/deletion when elements were added or removed from both W and C. We used the effective neutrality threshold approximation *N_e_*|*s*| *>* 1 to determine whether the effect of a mutation on fitness is strong enough to escape genetic drift. Considering that *Ne* ≃ *N* = 1, 000, mutations were categorized in their 3 categories: neutral (−0.001 *<* Δ*w <* 0.001), deleterious (Δ*w <* −0.001) and beneficial (Δ*w >* 0.001). Beneficial and deleterious were further split into 3 categories each: weak (|Δ*w*| *>* 0.001), mild (|Δ*w*| *>* 0.01), or strong (|Δ*w*| *>* 0.1) effects, to account for a total of 7 categories.

### Pleiotropy

Mutations can affect many, one, or none of the gene expressions. To determine the threshold from which we considered that the mutation affects gene expression, we considered the minimal amount of gene expression change Δ*P* that could trigger a non-neutral fitness change, *i.e.*, |Δ*w*| *>* 0.001 (our fitness effect threshold). The marginal bell-shaped fitness function on a single gene was 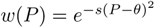, and the maximal fitness change was 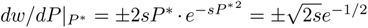 when 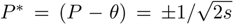 (the inflection points of the fitness function). For our default selection coefficient *s* = 10, this leads to a maximum effect on fitness of *dw/dP* ≃ 2.17. The gene expression change threshold from which Δ*w >* 0.001 was thus fixed at Δ*P* ≃ 0.001*/*2.17 = 0.00037. The pleiotropy level for a mutation was computed as the number of genes which expression change was ≥ Δ*P*.

### *Cis*- and *trans*-acting effects

*Cis*- and *trans*-effects were approximated by using the probability that a mutation impacts the expression of the very gene that received the mutation. If the gene harboring the mutation was among those with altered expression, the mutation was classified as *cis*-acting for that gene and *trans*-acting for all others; yet, the mutation is counted as having a *cis*-effect. If the gene harboring the mutation was *not* among those with altered expression, the mutation was considered to have *trans*-acting effects only. Using these measures, a probability of causing a *cis*-acting effect was computed for each level of pleiotropy.

## Results

In our individual-based simulations, the genotype of diploid, sexually-reproducing individuals encodes the structure of a gene regulatory network through two variables: a matrix **W** which elements quantifies the effect of each TF on the expression other genes, and a vector **C**, which elements stand for the activity of TFs. Simulations were run for 1,000 generations in a first environment (featured by a set of expression optima ***θ***_0_), and for 1,000 additional generations in a second environment representing a new adaptive challenge (Supplementary Figure S1). The potential beneficial or deleterious consequences of all kinds of mutations (regulatory, coding, gene duplications and deletions) was measured by simulating their fitness effects on genotypes sampled from generation 1,000 (adapted to the first environment) and assessed in the new environment.

### Initial Genotype and Gene Network Topology

Studying the effect of mutations on gene expression requires that models of gene networks reflect some realistic properties of biological systems. Yet, building a realistic structure for simulated networks is no trivial task. In particular, there is no guarantee that applying stabilizing selection on a subset of genes would necessarily lead to a realistic organization of gene interactions, as the evolved network structure might depend on the initial state of the system. We considered three network initialization methods at generation 0, leading to three evolutionary scenarios. In the ”full-active” scenario, all connections were initialized to random positive or negative values *W_ij_* ~ Gaussian(0, sd = 0.5) (hence ”full”), and all transcription factors were functional (hence ”active” : their activity was set to *C_i_* = 1) (Fig. 2A). In the ”empty-active” scenario, genes were not connected (*W_ij_* = 0) but they were functional (*C_i_* = 1). At least one mutation per gene was thus necessary to reach an expression level different from the constitutive expression (Fig. 2F). Finally, in the ”empty-inactive” scenario, genes were not connected (*W_ij_* = 0) and totally inactive (*C_i_* = 0). In these conditions, at least two mutations were necessary to change the expression of a gene: a mutation activating a transcription factor, and a mutation allowing the transcription factor to modify the expression of the target gene (Fig. 2K).

**Figure 2:**
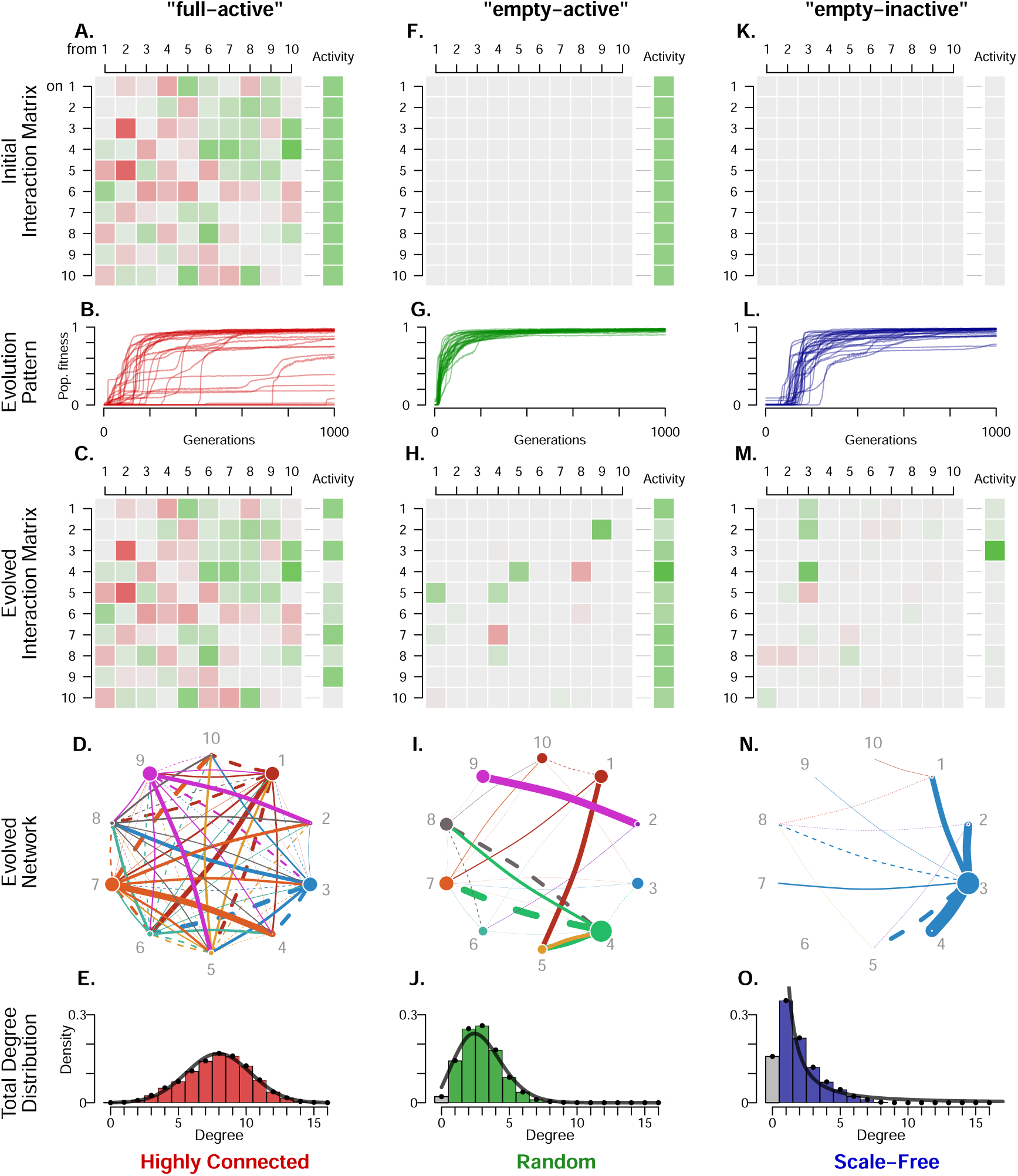
Three initialization scenarios evolved into distinct network topologies. **A-F-K**. Initial genotypes in the three scenarios, with elements of the regulatory interaction matrix initialized randomly (”full”) or set to zero (”empty”), and coding sequence activity set to 1 (”active” - green) or to zero (”inactive” - grey). The sign of the interaction is represented as red (repression) or green (activation), color intensity being proportional to the regulation strength. Adaptive evolution from these initial states was simulated for 1,000 generations, leading to an increase in population mean fitness (**B-G-L** - n=50 replicates shown). **C-H-M**. Examples of interaction matrices **W** after 1,000 generations, and their corresponding network graphical representation (**D-I-N**), where each gene is represented as one color. Genes 1 to 5 were submitted to selection for a random expression optimum, genes 6 to 10 were free to evolve. Edge color corresponds to the gene causing the regulation; activation: solid edges, repression: dashed. Node size is proportional to the activity *C_i_*, and edge width to the magnitude of the regulation *W_ij_*. **E-J-O**. Network total degree distributions at the end of the simulations (1,000 simulation replicates per topology) compared with Gaussian, Poisson ower law distributions, respectively.

Gene networks initiated using these three methods (1,000 replicates per method) were submitted to selection on gene expression for 5 out of 10 genes for 1,000 generations in populations of *N* =1,000 individuals: all initialized values of the genotype were thus allowed to change during the evolution. Those networks differed in terms of adaptation speed and maximum fitness achieved. The resulting adapted networks (Fig. 2C,H,M and 2D,I,N) illustrate different adaptation strategies. In the “full-active” scenario, the complex network connections did not change much, while TF activities evolved substantially. In the “empty-active” scenario, activities were modified to a lesser extent and new regulatory mutation had to evolve to connect the genes. In the “empty-inactive” scenario, only one gene (two in some simulation replicates) took the role of a ”hub” transcription factor that controls the expression of many other genes. Detailed network dynamics and alternative examples are available in Supplementary Movies S1 and S2.

The structure of gene regulatory networks was summarized by their degree distributions. While “full-active” and “empty-active” scenarios tended to generate Poisson degree distributions (with an average of 8 connections for “full-active”, and 3 connections for “empty-active”, Fig 2E and J), the degree distribution was close to a power law for the “empty-inactive” scenario, with an average of 2 connections per node (Fig 2O). These distributions match the network representations depicted in Fig 2D,I,N, each corresponding to a characteristic gene network topology: highly-connected networks, random networks (both Erdőos–Rényi-like), and scale-free networks (consistently obtained across varying parameters - see Supplementary Figure S2). These three distinct topologies form the basis for the analysis throughout this article.

Networks initiated as “empty-active” were the fastest to reach the gene expression optimum. Simulations were homogeneous, and fitness increased consistently with time. These networks also reached the highest average fitness (Fig. 2G). Networks from the “empty-inactive” scenario were slower to evolve (at least two mutations were required before any change in gene expression). The average final fitness was close to the “empty-active” adapted networks (Fig. 2L). In contrast, the average fitness of “full-active” networks was lower, and fitness gains seemed irregular and unpredictable (Fig. 2B). In some simulation replicates, the final fitness did not evolve much, indicating adaptation failures. Only 32% of the simulations starting from the “full-active” scenario were above 95% of the optimal fitness, versus 67% for “empty-inactive”, and 90% for “empty-active” simulations (see also Supplementary Figure S3). To make sure that functionally-equivalent networks were compared, this analysis (as well as the rest of the study) was performed from a subset of successful simulations, *i.e.*, that reached a population fitness of at least 0.95.

Network evolvabilities were then assessed by submitting successfully-evolved populations to a new evolutionary challenge, consisting in shifting randomly the optimal expression of 2 out of 5 genes, akin to a change from environment 1 to environment 2 (Fig. 3 and Supplementary Movies S1 and S2). The size of the shift was set such that the fitness dropped by about 50%. Networks were then submitted to stabilizing selection for the new optima for 1,000 additional generations. This second evolutionary stage consistently led to fast re-adaptation, and the relative performance of the three network topologies were more homogeneous than during the initial adaptation to the first environment. Random networks reached the optimal fitness slightly faster than scale-free networks, and highly-connected networks were slightly slower in average, although individual simulation runs largely overlapped. These results confirm that all three network topologies were able to produce potentially-adaptive mutations at a comparable (but not identical) rate.

**Figure 3:**
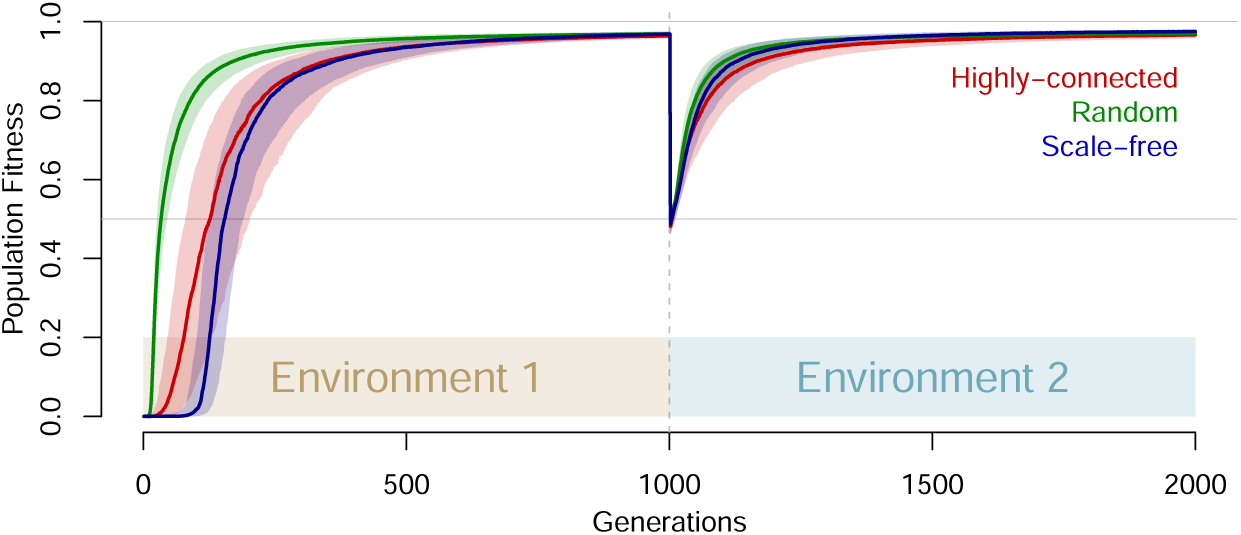
Three initial conditions and topologies drive adaptation dynamics. Solid lines are median of mean fitness across n=1,000 successful simulation replicates. Colored areas stand for the first and third quartiles of the distributions. The figure displays the first adaptation stage from an arbitrary genotypes starting point, and the second adaptation stage from an already-adapted network.

### Mutation Effects on Fitness

Our approach to study the effect of various mutation types on network adaptation was based on the ”re-adaptation” scenario from Fig. 3. Networks with different topologies were constructed from the three initial conditions described in Fig. 2, and their evolution was simulated for 1,000 generations in environment 1. Two gene expression optima were then changed randomly, and the fitness effect (Δ*w*, see Methods) of a series of random mutants in this new environment was calculated. The adaptation challenge was substantial, as fitness drops in average by 50% in environment 2. Three types of mutations were tested: mutations could target regulatory regions (affecting random elements in the matrix **W**), coding regions (affecting random genes in the vector **C**), or could change the number of genes (duplication of a random gene, or deletion of a non-selected gene). We characterized how genetic changes (mutation sizes - genotype) translate into gene expression (mutation effects - phenotype) and their consequences on fitness (fitness effects).

The distribution of the fitness effects of new mutations followed some general trends matching intuitive expectations (Fig. 4): (i) deleterious mutations were more frequent than beneficial mutations; (ii) deleterious mutations were often highly deleterious, beneficial mutations were often slightly beneficial; (iii) deleterious effects increased with the mutation size, but not beneficial effects, (iv) the lesser the effect of a mutation, the higher its probability of being selectively neutral. Other general observations, perhaps less intuitive, could also be described: (i) regulatory mutations, coding mutations, and deletion/duplications all have the capacity to participate to adaptation (although not to the same extent), and (ii) the distribution of fitness effects was roughly symmetrical for mutations that activate/inhibit gene expressions, decreased/increased gene activity, and delete/duplicate network genes.

**Figure 4:**
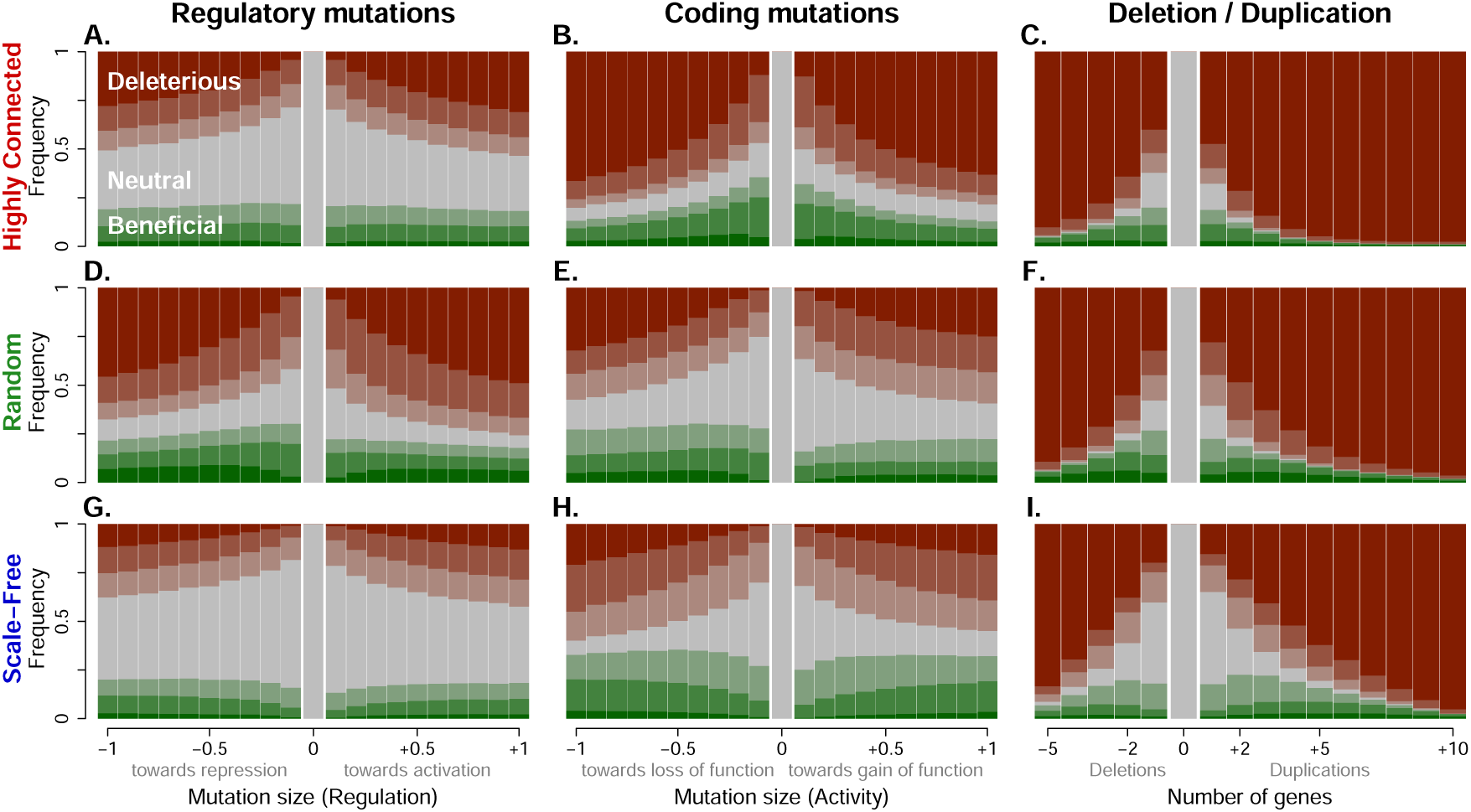
Fitness effects caused by different mutation types and sizes are influenced by network topology. The distribution of fitness effects Δ*w* was calculated in a putative evolutionary scenario where a population well-adapted to environment 1 (average fitness ≥ 0.95) was confronted to a new environment. At this point, single regulatory, coding, and deletion/duplication mutations of different arbitrary sizes (x-axis) were tested on 3 different network topologies (n=1,000 networks × 20 tests = 20,000 for each bar), resulting in either neutral (grey), beneficial (green) or deleterious (red) fitness effect of different magnitude (intensity of the color). The neutrality threshold was set to Δ*w <* 1*/N* = 0.001, and color intensities (red for deleterious mutations, green for advantageous mutations) correspond to |Δ*w*| *>* 0.001, 0.01, and 0.1 - weak, mild and strong effects respectively. Deletions and duplications were represented by the number of deleted (negative x-values) or duplicated (positive x-values) genes.

The distribution of fitness effects was heavily influenced by the network topology and by the nature of the mutations. Highly-connected networks were tolerant to regulatory mutations, but particularly sensitive to coding mutations. The opposite pattern was found for random networks, which were more sensitive to regulatory mutations and more tolerant to coding mutations. Scale-free networks appeared generally more tolerant to mutations (including gene deletions and duplications). Overall, there was no general adaptive advantage for a mutation type across all gene network topologies, except for deletions and duplications, which were more deleterious and less beneficial in all conditions.

### Pleiotropy and *cis*/*trans* Effects

The deleterious consequences of large-effect mutations are traditionally attributed to their pleiotropic effect, *i.e.*, their propensity to affect several traits at the same time. To explore the relationship between pleiotropy, mutation types, fitness effects, and network topology, we again carried out a series of mutations on adapted networks placed in a new environment. For regulatory and coding mutations, mutations of arbitrary size (−0.5 and +0.5) are shown here. For network size alterations, only single duplications are shown (because decreasing the network size mechanically decreases pleiotropy). The pleiotropic effect for each mutation was assessed by quantifying the number of genes which expression changed by more than a fixed value Δ*P* (see Methods). The fitness effect of each mutation (beneficial, neutral, or deleterious) was also recorded, along with its *cis*-effect, approximated as whether or not the gene that received the mutation had its own expression changed.

Virtually all regulatory mutations that changed gene expression in the network had *cis*-effects (Fig. 5): mutations directly target the gene regulation, and thus the gene expression. The mutated gene could then affect the rest of the network through cascading regulations, and thus the mutation display *trans*-effects in addition to a *cis*-effect. Note that regulatory mutations did not all have an effect on the network: some were silent (no gene affected), either because the mutation was compensated by feedbacks, or because the mutation changed the sensitivity to a transcription factor that was non-functional or not expressed. In contrast, virtually none of the coding mutations had *cis*-effects, unless they were very pleiotropic — when the whole network was disturbed, the gene carrying the coding mutation was necessarily affected as well through cascading effects looping back to the original gene.

**Figure 5:**
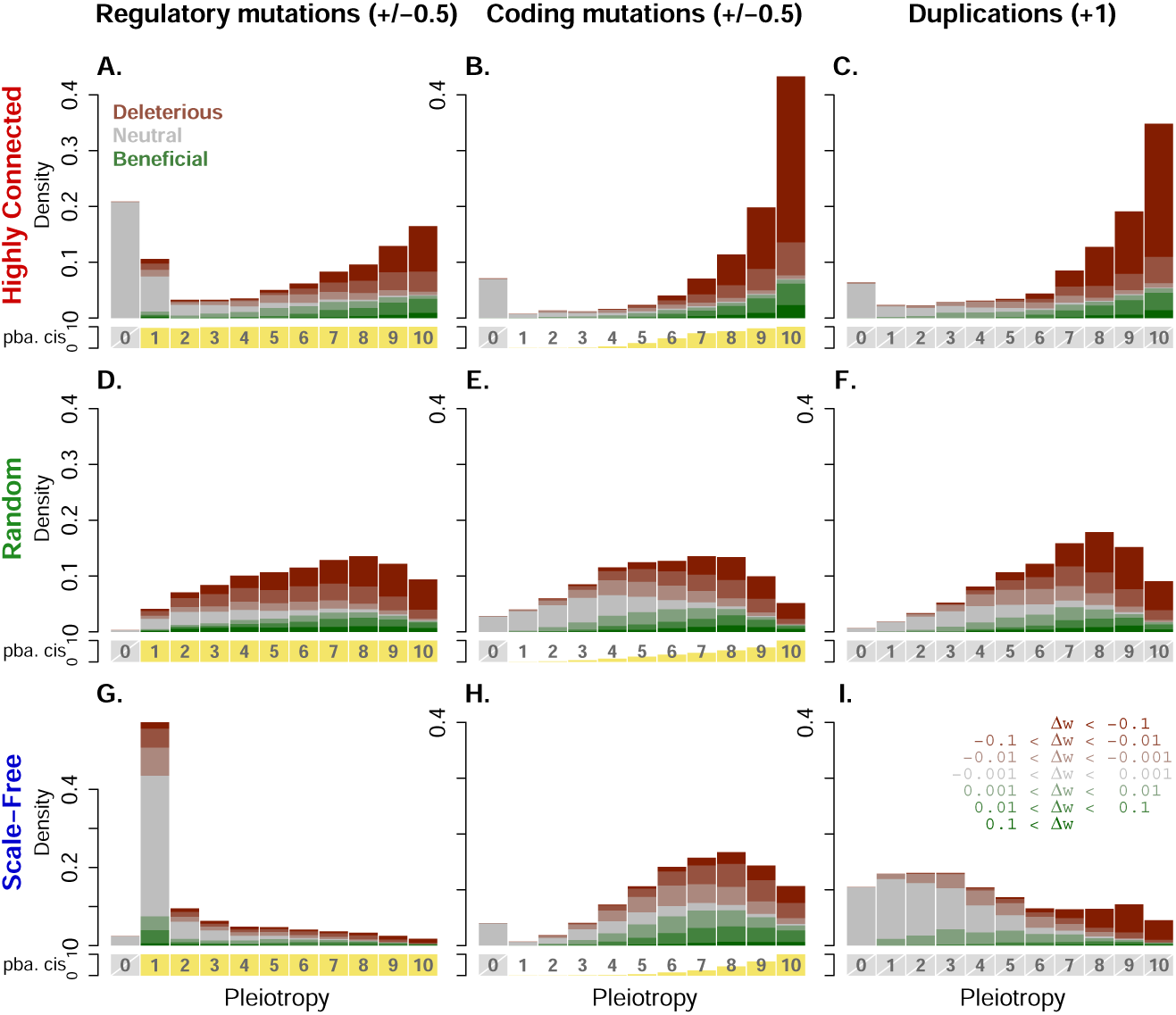
Pleiotropic and *cis*-acting effects of different mutation types are influenced by network topology. Pleiotropy in the x-axis, was computed as the number of genes in the network which expression was altered by the mutation. Lower panels in yellow: probability of a mutation to trigger a *cis*-effect (not applicable for pleiotropy of 0 or duplications, hence their grey status); main panels: pleiotropy density distribution as fractions of fitness effects (deleterious, neutral and beneficial, in red, grey and green respectively - color intensity according to changes in fitness Δ*w*’s intervals). For each of the three topologies and three mutation types, *n* = 50, 000 mutations were carried out upon environmental change: regulatory and coding mutations were set to ±0.5 and duplications to +1.

Here again, network topology largely influenced the distribution of pleiotropic effects. Highly-connected networks were characterized by a U-shape distribution of pleiotropic effects, with two categories of mutations: no-effect neutral mutations, and large-effect, mostly deleterious mutations. All three mutation types (regulatory, coding, and duplications) displayed the same pattern. In contrast, pleiotropy in random networks had a single, intermediate mode; most mutations affected between 3 and 9 genes, irrespective for the mutation type. Highly-pleiotropic mutations were very deleterious, low-pleiotropy mutations were rather neutral. The pleiotropic pattern for scale-free networks was more complicated, and differed according to the mutation type. While coding mutations were similar to random networks (intermediate mode), regulatory mutations were featured by a very frequent ”single gene effect” category, concentrating most neutral and beneficial mutations. The distribution of pleiotropy in duplications was also specific, both less pleiotropic and less deleterious than in the other networks.

All three network topologies displayed some form of robustness to mutations, although the mechanisms by which mutational effects were buffered differed substantially. In highly-connected networks, a substantial fraction of mutations had no effect on gene expression (pleiotropy = 0). These ”silent” mutations were thus necessarily selectively neutral, but non-silent mutations affected multiple genes and had large fitness consequences. In contrast, random networks were virtually deprived of silent mutations. Yet, the effect of coding mutations on fitness was smaller and more neutral than for other networks. Buffering in scale-free networks was also different: most regulatory mutations were not pleiotropic, and had thus a higher chance to affect only non-selected genes.

Overall, the more a mutation was pleiotropic, the larger its effect on fitness (see also Supplementary Figure S4). Larger mutations resulted in greater pleiotropic effects, while the overall distribution remained similar (Supplementary Figure S5). Mutations on selected and non-selected (free) genes had very similar pleiotropic effects, but they necessarily differed in the size of their fitness consequences (Supplementary Figure S6). This aligns with the expectation that pleiotropic mutations are generally more deleterious. Yet, in many cases, the proportion of beneficial mutations increased with pleiotropy: strongly-beneficial mutations were also found at the highest pleiotropy degree, together with the most deleterious mutations. This suggests that pleiotropy might be both highly deleterious and frequently involved in quick adaptation, which can appear paradoxical but not contradictory.

Finally, we determined the properties of the mutations that were expected to participate preferentially to adaptation from a large set of coding and regulatory mutations of various sizes. In scale-free networks, beneficial mutations were enriched in coding, pleiotropic, *trans*- and large-effect mutations (figure 6). In contrast, most neutral mutations were regulatory, with low pleiotropy. The properties of deleterious vs. beneficial mutations were rather similar, deleterious mutations being slightly more pleiotropic than beneficial mutations. In highly-connected and random networks, pleiotropic mutations were also more represented among beneficial mutations (Supplementary Figure S7). However, here again, the mutation type preference was topology-dependent: in highly-connected networks, beneficial (and deleterious) mutations were enriched for coding and *cis*-acting mutations, while beneficial (and deleterious) mutations in random networks involved regulatory and *cis*-acting mutations. Overall, (i) beneficial mutations were more similar to deleterious mutations than to neutral mutations, and (ii) apart from a larger degree of pleiotropy, shared over all genetic architectures, the enrichment of beneficial mutations in *cis*- vs. *trans*-effects, in regulatory vs. coding effects, and in large vs. small molecular effect depends on the topology of the gene regulatory network.

**Figure 6:**
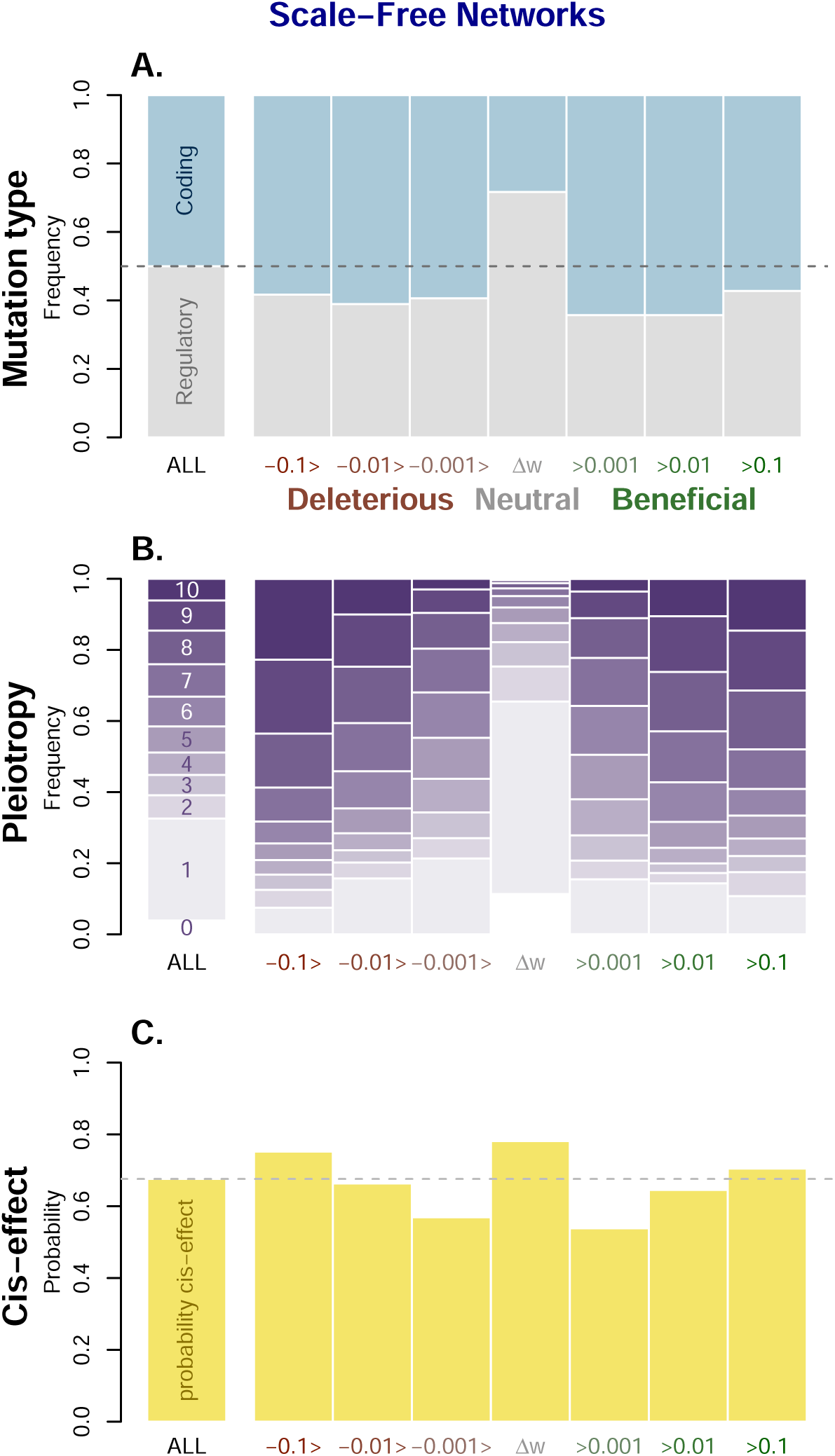
Mutation properties as a function of Fitness effects in scale-free networks. *n* = 100, 000 mutations (from 1, 000 independent networks) were generated and classified according to changes in fitness Δ*w*’s intervals. Y-axis: proportions for (**A**.) regulatory vs. coding mutations, (**B**.) pleiotropy, and (**C**.) the probability of a mutation to trigger a *cis*-effect.

## Discussion

Coupling a gene regulatory model with a population genetics framework makes a powerful theoretical tool to simulate the fitness effects of various types of mutations affecting gene expression. We manipulated the gene network topology by modifying the initial genotype, and we surveyed the evolutionary dynamics of gene networks in simulated populations submitted to an adaptive scenario. Simulations showed that the speed of adaptation (and thus, the rate and size of beneficial mutations) depends on the gene network topology. Moreover, the distribution of fitness effects of different types of mutations was also heavily affected by the structure of the network, and the consequences of regulatory mutations, alterations of the coding sequence of transcription factors, and gene duplications and deletions was substantially different in random, highly-connected, and scale-free networks. As explanatory mechanism, we suggest that pleiotropy, by reflecting topological properties, constrains the effects of mutations. Thus we found that the widely-accepted idea that regulatory mutations were less deleterious and less pleiotropic than mutations in the coding sequence was only valid in scale-free regulatory networks. The evolutionary potential of regulatory, coding, or duplication/deletion mutations may indeed differ, but this variation reflects the organization of the regulatory system rather than intrinsic mutational properties.

### Gene Regulation Model

Our model implements one of many existing theoretical abstractions of regulation mechanisms used to analyze the properties and evolution of gene regulatory networks. The simplest models generally rely on Boolean networks (Kauffman 1969; Mayer and Hansen 2017), while more realistic but more computationally-intensive models are based on systems of non-linear differential equations derived from biochemistry and thermodynamics theory (Polynikis et al. 2009). Our approach builds on Wagner’s model (A. Wagner 1994; A. Wagner 1996), which offers intermediate complexity with simple (additive) but explicit regulatory interactions. This popular model enables the study of both the quantitative evolution of gene expression regulation and the topology of the network (Azevedo et al. 2006; Bergman and Siegal 2003; Ciliberti et al. 2007; Fierst and Phillips 2015).

Our implementation of the model differed from the original one by the scaling function (Equation 2, as in e.g. Rünneburger and Le Rouzic (2016)) which ensures (i) a quantitative measurement of gene expression scaled between 0 and 1, (ii) a constitutive expression that differs from the mid-point between minimum and maximum expression. Specifically for this study, we introduced coding mutations through an explicit genetic vector (**C**), representing the regulatory activity of the TF proteins.

Every model relies on simplifications, and some features of the simulated genetic architectures remain far from the complexity of real regulatory mechanisms. For instance, the probabilities and sizes of gain-of-function versus loss-of-function for coding mutations were identical, which is not typical for protein-coding genes (Gerasimavicius et al. 2022). Symmetrical activation/inhibition rates are probably less unrealistic for transcription factor binding sites, since (i) binding sites often appear spontaneously in random sequences, making balanced gain/loss of function mutations more probable (Berg et al. 2004; Doniger and Fay 2007; Tuğrul et al. 2015), and (ii) many transcription factors act as both repressors and activators for different genes (Bauer et al. 2010; Boyle and Després 2010).

### Properties of Biological Networks

Our results highlight the essential role of gene network topology on the pleiotropy and fitness effect distribution of new mutations. The three gene network topologies we explored in our simulations were not pre-determined by arbitrary connection rules, but rather emerged from different initial genotypes in the simulations. These genotypes then evolved under the same mutation-selection process. As a consequence, we could only study general network topologies achievable from a restricted set of initial conditions, leaving more precise topological features unexplored, such as hierarchical patterns in scale-free networks (Ravasz and Barabási 2003).

Here, we applied two modes of selection, stabilizing selection (penalizing genotypes which target genes deviate from the optimal expression), and stability selection (fitness penalty for cycling networks). Both stabilizing and stability selection are known to promote network robustness through indirect selection: stabilizing selection slightly penalizes genotypes which mutant offspring deviates strongly from the optimum (Rünneburger and Le Rouzic 2016; G. P. Wagner, Booth, and Bagheri-Chaichian 1997), while the network propensity to cycle (instability) happens to be correlated with the sensitivity to disturbances (Le Rouzic 2022; Siegal and Bergman 2002). To date, the role of selection into the evolution of network robustness remains rather elusive, but some local topological properties, such as the enrichment in feed-forward regulatory loops (Cooper et al. 2008; Mangan et al. 2006), could realistically result from selection against expression noise (Ghosh et al. 2005; Osella et al. 2011).

In our simulations, some topologies were clearly suboptimal; highly-connected networks displayed poor adaptive properties, with all types of mutations being more pleiotropic and deleterious compared to other network topologies. While we did not quantify robustness and evolvability, our results appear coherent with other studies showing that scale-free networks harbor faster and smoother evolutionary trajectories and have greater robustness to mutational and environmental perturbations, compared to random networks (Greenbury et al. 2010; Oikonomou and Cluzel 2006).

Of particular interest is the emergence of scale-freeness in artificial and biological networks. Although it is still being debated (Smith et al. 2021), the scale-free (or ”scale-free-like”) topology is often considered as prevalent in biological networks, including in gene regulatory networks (Fagny and Austerlitz 2021; Wuchty et al. 2006). It has been proposed that the evolution of large-scale features, such as scale-freeness, might result from non-adaptive processes, including mutational bias and systems drift (M. Lynch 2007). Scale-freeness could indeed emerge from the intrinsic process of gene network evolution (growth by duplication and *de novo* genes expressed as soon as they connect to existing regulators (Yona et al. 2018)), suggesting that such a universality may not need any additional adaptive conjecture (Chung et al. 2003). In our simulations, we obtained a scale-free topology without constraining the network, but only by requiring independent mutations to turn on the regulatory effects and the activity of transcription factors. To some extent, this is equivalent of letting the network grow step by step, in line with the ”preferential attachment” mechanism.

Yet, even if regulatory networks tend to be scale-free at a large scale, local regulatory patterns may substantially deviate from the power-law distribution (Winterbach et al. 2013). The gene position in the network can influence its propensity to be a target for adaptive evolution across different mutation types, emphasizing the importance of local topology (Fagny and Austerlitz 2021). For example, Stern and Orgogozo (2009) argues that hotspot genes are more likely to undergo evolutionary change. It has also been shown that local regulatory structures reflect the biological function of the modules Luscombe et al. (2004), which may not be true at a larger scale. The 10-gene regulatory networks in our simulations, perhaps closer to actual regulatory ”motifs”, correspond to this local regulation level, and unusual connection patterns (including random or highly-connected networks) are not necessarily unrealistic at such a scale.

### Genetic Bases of Adaptation

Our results thus brought little evidence that regulatory changes, coding mutations, or gene duplications or deletions may be intrinsically more prone to contribute to adaptation. Yet, differences in pleiotropy and fitness effects were substantial, and these differences emerged from the structure of the regulatory network, with different topologies leading to varying sensitivities to different mutation types. Although not completely intuitive, we argue this can be reasonably explained. For instance, regulatory mutations in important transcription factors carry a substantial cascading potential, and high levels of pleiotropy. Similarly, the immediate consequences of gene duplications and deletions, which drastically change gene expression, may not be fundamentally different from large-effect regulatory mutations doubling or canceling gene expression. Thus, the contribution of different mutation types to the pool of potentially adaptive alleles may not be rooted in the molecular nature of the mutation but rather in subtle features of the biological system, including the topology of the regulatory network.

The Distribution of Fitness Effects (DFEs) of new mutations has received some substantial theoretical and empirical interest, as it conditions the evolutionary properties of populations (Eyre-Walker and Keightley 2007). In the last decades, progress in genomics and population genetics made it possible to identify the molecular nature of individual mutations, and thus to estimate specific DFEs in model organisms, offering an opportunity for comparison with our simulation results. Contrasting regulatory and coding mutations segregating in human populations supports larger effect of coding mutations on fitness (Racimo and Schraiber 2014), which is consistent with our scale-free network simulations. In primates, most non-synonymous coding mutations have a mild deleterious effect, less than 15% being very deleterious (Eyre-Walker, Woolfit, and Phelps 2006), and up to 15% being beneficial (Castellano et al. 2019). In yeast, *trans*-acting regulatory mutations are common, but they are more pleiotropic and more deleterious than *cis*-acting regulatory mutations (Schaefke et al. 2013; Vande Zande, Hill, and Wittkopp 2022; Wittkopp, Haerum, and Clark 2008). This pleiotropic burden of *trans*-acting mutations was consistent with our scale-free network simulations, but not with other network topologies. Notably, this is an observation that has recently received some empirical support (Vande Zande and Wittkopp 2022). Gene duplications and deletions were proven to affect fitness to a larger extent than other types of mutations in a large variety of organisms, including *E. coli* (Adler et al. 2014), yeast (Payen et al. 2016), and drosophila (Emerson et al. 2008). Duplications tend to be less deleterious than deletions, and more prone to be recruited during adaptation (Emerson et al. 2008; Payen et al. 2016), which was again predicted in our scale-free network simulations.

The literature frequently suggests that coding mutations, being more pleiotropic, are more deleterious and thus less likely to be selected for adaptation (Carroll 2005; Wittkopp, Haerum, and Clark 2008). Although our simulations confirm that coding mutations are indeed more often deleterious than regulatory mutations in scale-free networks, they also appear to be more beneficial in our evolutionary scenario. It is difficult to be conclusive on this issue, as empirical DFEs rarely estimate both deleterious and beneficial mutations together: mutation-accumulation experiments generally generate neutral and deleterious mutations (Bao et al. 2022), experimental evolution accumulates adaptive mutations (MacLean and Buckling 2009), and methods based on the frequency spectrum of segregating alleles are rarely able to identify ancestral vs. derived alleles (but see Castellano et al. (2019) for an exception). In any case, our arbitrary evolutionary scenario implied the recovery from a large (~ 50%) fitness drop, which may be out of the range of typical adaptive evolution, and may favor unrealistic large-effect mutations. Whether or not large and pleiotropic coding mutations can be retained during adaptation thus remains an open question.

Our theoretical model confirmed that all types of mutations (regulatory, coding, duplications and deletions) have the potential to contribute to adaptive evolution of gene expression. This echoes accumulating observations about the diversity of the molecular bases of adaptation. Yet, this does not mean that all types of mutations must be equally represented among beneficial alleles, as any predominant form of adaptive mutation could realistically emerge as a consequence of differences in, e.g., mutation rates, rather than from intrinsic advantages to specific molecular mechanisms. It is also likely that traits considered at different organismal levels (gene expression and metabolism vs. morphology or behavior) may differ in terms of genetic architectures, and thus in the distribution of fitness effects of mutations. We believe that theoretical approaches offer alternative ways to confirm and challenge our understanding of evolutionary systems biology, and feed transcriptomics, comparative genomics, or synthetic biology that can then be tested empirically.

## Acknowledgements

We thank Apolline Petit and Fabien Duveau for their thorough reading of the manuscript and their valuable suggestions. We also warmly thank Pascal Hersen for allowing us to perform preliminary bioinformatics simulations and analyses at the Institut Curie in Paris.

## Data availability

The simulation program and the analysis scripts were written in R version 4.04 (R Core Team 2021), and are available as Supplementary Material and in the dedicated GitHub repository: https://github.com/spouze/GeneRation_Pouzet2024/.

## Author contributions

S. Pouzet: Software development, Data analysis, Writing. A. Le Rouzic: Supervision, Writing.

## References

Adler, M., Anjum, M., Berg, O. G., Andersson, D. I., & Sandegren, L. (2014). High fitness costs and instability of gene duplications reduce rates of evolution of new genes by duplication-divergence mechanisms. Molecular biology and evolution, 31 (6), 1526–1535.

Aguilar-Rodríguez, J., & Payne, J. L. (2021). Robustness and evolvability in transcriptional regulation. *Evolutionary Systems Biology: Advances*, Questions, and Opportunities, 197–219. 10.1007/978-3-030-71737-79

Azevedo, R. B., Lohaus, R., Srinivasan, S., Dang, K. K., & Burch, C. L. (2006). Sexual reproduction selects for robustness and negative epistasis in artificial gene networks. Nature, 440 (7080), 87–90.

Bao, K., Melde, R. H., & Sharp, N. P. (2022). Are mutations usually deleterious? a perspective on the fitness effects of mutation accumulation. Evolutionary ecology, 36 (5), 753–766.

Bauer, D. C., Buske, F. A., & Bailey, T. L. (2010). Dual-functioning transcription factors in the developmental gene network of drosophila melanogaster. BMC bioinformatics, 11, 1–14. 10.1186/1471-2105-11-366

Berg, J., Willmann, S., & Lässig, M. (2004). Adaptive evolution of transcription factor binding sites. BMC evolutionary biology, 4, 1–12. 10.1186/1471-2148-4-42

Bergman, A., & Siegal, M. L. (2003). Evolutionary capacitance as a general feature of complex gene networks. Nature, 424 (6948), 549–552.

Boyle, P., & Després, C. (2010). Dual-function transcription factors and their entourage: Unique and unifying themes governing two pathogenesis-related genes. Plant signaling & behavior, 5 (6), 629–634. 10.4161/psb.5.6.11570

Carroll, S. B. (2005). Evolution at two levels: On genes and form. PLoS biology, 3 (7), e245. 10.1371/journal.pbio.0030245

Carroll, S. B. (2008). Evo-devo and an expanding evolutionary synthesis: A genetic theory of morphological evolution. Cell, 134 (1), 25–36. 10.1016/j.cell.2008.06.030

Castellano, D., Macià, M. C., Tataru, P., Bataillon, T., & Munch, K. (2019). Comparison of the full distribution of fitness effects of new amino acid mutations across great apes. Genetics, 213 (3), 953–966.

Chung, F., Lu, L., & Vu, V. (2003). Spectra of random graphs with given expected degrees. Proceedings of the National Academy of Sciences of the United States of America, 100 (11), 6313–6318. 10.1073/pnas.0937490100

Ciliberti, S., Martin, O. C., & Wagner, A. (2007). Innovation and robustness in complex regulatory gene networks. Proceedings of the National Academy of Sciences, 104 (34), 13591–13596.

Coolon, J. D., McManus, C. J., Stevenson, K. R., Graveley, B. R., & Wittkopp, P. J. (2014). Tempo and mode of regulatory evolution in drosophila. Genome research, 24 (5), 797–808.

Cooper, M. B., Loose, M., & Brookfield, J. F. (2008). Evolutionary modelling of feed forward loops in gene regulatory networks. BioSystems, 91 (1), 231–244. 10.1016/j.biosystems.2007.09.004

Crombach, A., & Hogeweg, P. (2008). Evolution of evolvability in gene regulatory networks. PLoS computational biology, 4 (7), e1000112.

Doniger, S. W., & Fay, J. C. (2007). Frequent gain and loss of functional transcription factor binding sites. PLoS Computational Biology, 3 (5), 0932–0942. 10.1371/ journal.pcbi.0030099

Emerson, J., Cardoso-Moreira, M., Borevitz, J. O., & Long, M. (2008). Natural selection shapes genome-wide patterns of copy-number polymorphism in *Drosophila melanogaster*. Science, 320 (5883), 1629–1631.

Eyre-Walker, A., & Keightley, P. D. (2007). The distribution of fitness effects of new mutations. Nature Reviews Genetics, 8 (8), 610–618.

Eyre-Walker, A., Woolfit, M., & Phelps, T. (2006). The distribution of fitness effects of new deleterious amino acid mutations in humans. Genetics, 173 (2), 891–900.

Fagny, M., & Austerlitz, F. (2021). Polygenic adaptation: Integrating population genetics and gene regulatory networks. Trends in Genetics, 37 (7), 631–638.

Fierst, J. L., & Phillips, P. C. (2015). Modeling the evolution of complex genetic systems: The gene network family tree. Journal of Experimental Zoology Part B: Molecular and Developmental Evolution, 324 (1), 1–12. 10.1002/jez.b.22597

Fraser, H. B. (2013). Gene expression drives local adaptation in humans. Genome Research, 23 (7), 1089–1096. 10.1101/gr.152710.112

Galant, R., & Carroll, S. B. (2002). Evolution of a transcriptional repression domain in an insect hox protein. Nature, 415 (6874), 910–913. 10.1038/nature717

Gerasimavicius, L., Livesey, B. J., & Marsh, J. A. (2022). Loss-of-function, gain-of-function and dominant-negative mutations have profoundly different effects on protein structure. Nature Communications, 13 (1), 1–15. 10.1038/s41467-022-31686-6

Ghosh, B., Karmakar, R., & Bose, I. (2005). Noise characteristics of feed forward loops. Physical Biology, 2 (1), 36–45. 10.1088/1478-3967/2/1/005

Greenbury, S. F., Johnston, I. G., Smith, M. A., Doye, J. P., & Louis, A. A. (2010). The effect of scale-free topology on the robustness and evolvability of genetic regulatory networks. Journal of theoretical biology, 267 (1), 48–61.

Hoekstra, H. E., & Coyne, J. A. (2007). The locus of evolution: Evo-Devo and the genetics of adaptation. Evolution, 61 (5), 995–1016. 10.1111/j.1558-5646.2007.00105.x

Kaneko, T., & Kikuchi, M. (2022). Evolution enhances mutational robustness and suppresses the emergence of a new phenotype: A new computational approach for studying evolution. PLOS Computational Biology, 18 (1), e1009796.

Kauffman, S. A. (1969). Metabolic stability and epigenesis in randomly constructed genetic nets. Journal of theoretical biology, 22 (3), 437–467. 10.1016/0022-5193(69)90015-0

King, M.-C., & Wilson, A. C. (1975). Evolution at two levels in humans and chimpanzees: Their macromolecules are so alike that regulatory mutations may account for their biological differences. Science, 188 (4184), 107–116. 10.1126/science.1090005

Kondrashov, F. A. (2012). Gene duplication as a mechanism of genomic adaptation to a changing environment. Proceedings of the Royal Society B: Biological Sciences, 279 (1749), 5048– 5057. 10.1098/rspb.2012.1108

Krishnan, J., Seidel, C. W., Zhang, N., Singh, N. P., VanCampen, J., Peuß, R., Xiong, S., Kenzior, A., Li, H., Conaway, J. W., et al. (2022). Genome-wide analysis of cis-regulatory changes underlying metabolic adaptation of cavefish. Nature genetics, 54 (5), 684–693.

Kuo, P. D., Banzhaf, W., & Leier, A. (2006). Network topology and the evolution of dynamics in an artificial genetic regulatory network model created by whole genome duplication and divergence. Biosystems, 85 (3), 177–200.

Le Rouzic, A. (2022). Gene network robustness as a multivariate character. Peer Community Journal, 2, e26. 10.24072/pcjournal.125

López-Maury, L., Marguerat, S., & Bähler, J. (2008). Tuning gene expression to changing environments: from rapid responses to evolutionary adaptation. Nature Reviews Genetics, 9 (8), 583–593. 10.1038/nrg2398

Luscombe, N. M., Madan Babu, M., Yu, H., Snyder, M., Teichmann, S. A., & Gerstein, M. (2004). Genomic analysis of regulatory network dynamics reveals large topological changes. Nature, 431 (7006), 308–312. 10.1038/nature02782

Lynch, M. (2007). The evolution of genetic networks by non-adaptive processes. Nature Reviews Genetics, 8 (10), 803–813. 10.1038/nrg2192

MacLean, R. C., & Buckling, A. (2009). The distribution of fitness effects of beneficial mutations in pseudomonas aeruginosa. PLoS genetics, 5 (3), e1000406.

Mangan, S., Itzkovitz, S., Zaslaver, A., & Alon, U. (2006). The incoherent feed-forward loop accelerates the response-time of the gal system of Escherichia coli. Journal of Molecular Biology, 356 (5), 1073–1081. 10.1016/j.jmb.2005.12.003

Marand, A. P., Eveland, A. L., Kaufmann, K., & Springer, N. M. (2023). Cis-regulatory elements in plant development, adaptation, and evolution. Annual review of plant biology, 74 (1), 111–137.

Mayer, C., & Hansen, T. F. (2017). Evolvability and robustness: A paradox restored. Journal of theoretical biology, 430, 78–85. 10.1016/j.jtbi.2017.07.004

Moses, A. M., Pollard, D. A., Nix, D. A., Iyer, V. N., Li, X.-Y., Biggin, M. D., & Eisen, M. B. (2006). Large-scale turnover of functional transcription factor binding sites in drosophila. PLoS computational biology, 2 (10), e130. 10.1371/journal.pcbi.0020130

Oikonomou, P., & Cluzel, P. (2006). Effects of topology on network evolution. Nature Physics, 2 (8), 532–536.

Osella, M., Bosia, C., Corá, D., & Caselle, M. (2011). The role of incoherent microRNA-mediated feedforward loops in noise buffering. PLoS Computational Biology, 7 (3), e1001101. 10.1371/journal.pcbi.1001101

Pavlicev, M., Bourg, S., & Le Rouzic, A. (2022). The genotype-phenotype map structure and its role for evolvability.

Payen, C., Sunshine, A. B., Ong, G. T., Pogachar, J. L., Zhao, W., & Dunham, M. J. (2016). High-throughput identification of adaptive mutations in experimentally evolved yeast populations. PLoS genetics, 12 (10), e1006339.

Polynikis, A., Hogan, S., & Di Bernardo, M. (2009). Comparing different ode modelling approaches for gene regulatory networks. Journal of theoretical biology, 261 (4), 511–530. 10.1016/j.jtbi.2009.07.040

Prud’homme, B., Gompel, N., & Carroll, S. B. (2007). Emerging principles of regulatory evolution. Proceedings of the National Academy of Sciences, 104 (suppl 1), 8605–8612. 10.1073/pnas.0700488104

Qian, W., & Zhang, J. (2014). Genomic evidence for adaptation by gene duplication. Genome Research, 24 (8), 1356–1362. 10.1101/gr.172098.114

Racimo, F., & Schraiber, J. G. (2014). Approximation to the distribution of fitness effects across functional categories in human segregating polymorphisms. PLoS genetics, 10 (11), e1004697.

Ravasz, E., & Barabási, A.-L. (2003). Hierarchical organization in complex networks. Physical review E, 67 (2), 026112.

Ronshaugen, M., McGinnis, N., & McGinnis, W. (2002). Hox protein mutation and macroevolution of the insect body plan. Nature, 415 (6874), 914–917. 10.1038/nature716

Rünneburger, E., & Le Rouzic, A. (2016). Why and how genetic canalization evolves in gene regulatory networks. BMC evolutionary biology, 16, 1–11. 10.1186/s12862-016-0801-2

Sawyer, S. A., Kulathinal, R. J., Bustamante, C. D., & Hartl, D. L. (2003). Bayesian analysis suggests that most amino acid replacements in drosophila are driven by positive selection. Journal of molecular evolution, 57, S154–S164.

Schaefke, B., Emerson, J., Wang, T.-Y., Lu, M.-Y. J., Hsieh, L.-C., & Li, W.-H. (2013). Inheritance of gene expression level and selective constraints on *trans*-and *cis*-regulatory changes in yeast. Molecular biology and evolution, 30 (9), 2121–2133.

Siegal, M. L., & Bergman, A. (2002). Waddington’s canalization revisited: Developmental stability and evolution. Proceedings of the National Academy of Sciences, 99 (16), 10528–10532. 10.1073/pnas.102303999

Signor, S. A., & Nuzhdin, S. V. (2018). The evolution of gene expression in cis and trans. Trends in Genetics, 34 (7), 532–544.

Smith, H. B., Kim, H., & Walker, S. I. (2021). Scarcity of scale-free topology is universal across biochemical networks. Scientific reports, 11 (1), 6542. 10.1038/s41598-021-85903-1

Solé, R. V., & Valverde, S. (2008). Spontaneous emergence of modularity in cellular networks. Journal of The Royal Society Interface, 5 (18), 129–133.

Stern, D. L., & Orgogozo, V. (2008). The loci of evolution: How predictable is genetic evolution? Evolution, 62 (9), 2155–2177. 10.1111/j.1558-5646.2008.00450.x

Stern, D. L., & Orgogozo, V. (2009). Is genetic evolution predictable? Science, 323 (5915), 746–751. 10.1126/science.1158997

Teichmann, S. A., & Babu, M. M. (2004). Gene regulatory network growth by duplication. Nature genetics, 36 (5), 492–496. 10.1038/ng1340

Tuğrul, M., Paixao, T., Barton, N. H., & Tkačik, G. (2015). Dynamics of transcription factor binding site evolution. PLoS genetics, 11 (11), e1005639. 10.1371/journal.pgen.1005639

Vande Zande, P., Hill, M. S., & Wittkopp, P. J. (2022). Pleiotropic effects of trans-regulatory mutations on fitness and gene expression. Science, 377 (6601), 105–109.

Vande Zande, P., & Wittkopp, P. J. (2022). Network topology can explain differences in pleiotropy between cis-and trans-regulatory mutations. Molecular Biology and Evolution, 39 (12), msac266.

Wagner, A. (1994). Evolution of gene networks by gene duplications: A mathematical model and its implications on genome organization. Proceedings of the National Academy of Sciences, 91 (10), 4387–4391. 10.1073/pnas.91.10.4387

Wagner, A. (1996). Does evolutionary plasticity evolve? Evolution, 50 (3), 1008–1023. 10.1111/j.1558-5646.1996.tb02342.x

Wagner, G. P., Booth, G., & Bagheri-Chaichian, H. (1997). A population genetic theory of canalization. Evolution, 51 (2), 329–347. 10.1111/j.1558-5646.1997.tb02420.x

Wagner, G. P., & Lynch, V. J. (2008). The gene regulatory logic of transcription factor evolution. Trends in Ecology & Evolution, 23 (7), 377–385.

Winterbach, W., Mieghem, P. V., Reinders, M., Wang, H., & Ridder, D. D. (2013). Topology of molecular interaction networks. BMC systems biology, 7, 1–15. 10.1186/ 1752-0509-7-90

Wittkopp, P. J., Haerum, B. K., & Clark, A. G. (2008). Regulatory changes underlying expression differences within and between drosophila species. Nature genetics, 40 (3), 346–350.

Wittkopp, P. J., & Kalay, G. (2012). Cis-regulatory elements: Molecular mechanisms and evolutionary processes underlying divergence. Nature Reviews Genetics, 13 (1), 59–69.

Wray, G. A. (2007). The evolutionary significance of cis-regulatory mutations. Nature Reviews Genetics, 8 (3), 206–216.

Wuchty, S., Ravasz, E., & Barabási, A.-L. (2006). The architecture of biological networks. Complex systems science in biomedicine, 165–181.

Yona, A. H., Alm, E. J., & Gore, J. (2018). Random sequences rapidly evolve into de novo promoters. Nature communications, 9 (1), 1530. 10.1038/s41467-018-04026-w

Zhang, J. (2003). Evolution by gene duplication: An update. Trends in ecology & evolution, 18 (6), 292–298. 10.1016/S0169-5347(03)00033-8

