## Supplementary Material for "Gene network topology drives the mutational landscape of gene expression"

**Data availability**: The simulation program and the analysis scripts were written in R version 4.04 (R Core Team 2021), and are available as Supplementary Material and in the dedicated GitHub repository:
`https://github.com/spouze/GeneRation_Pouzet2024/`.

**Author contributions**: S. Pouzet: Software development, Data analysis, Writing. A. Le Rouzic: Supervision, Writing.

**Funding and support**: The main simulations were carried out on the Core Cluster of the Institut Français de Bioinformatique (IFB) (ANR-11-INBS-0013). The authors acknowledge the ANR – FRANCE (French National Research Agency) for its financial support of the Evoplanet project n°ANR-22-CE02-0026.

**Conflict of interest statement**: the authors declare no conflict of interest.

**Acknowledgements**: We thank Apolline Petit and Fabien Duveau for their thorough reading of the manuscript and their valuable suggestions. We also warmly thank Pascal Hersen for allowing us to perform preliminary bioinformatics simulations and analyses at the Institut Curie in Paris.

### Supplementary material for: Gene network topology drives the mutational landscape of gene expression

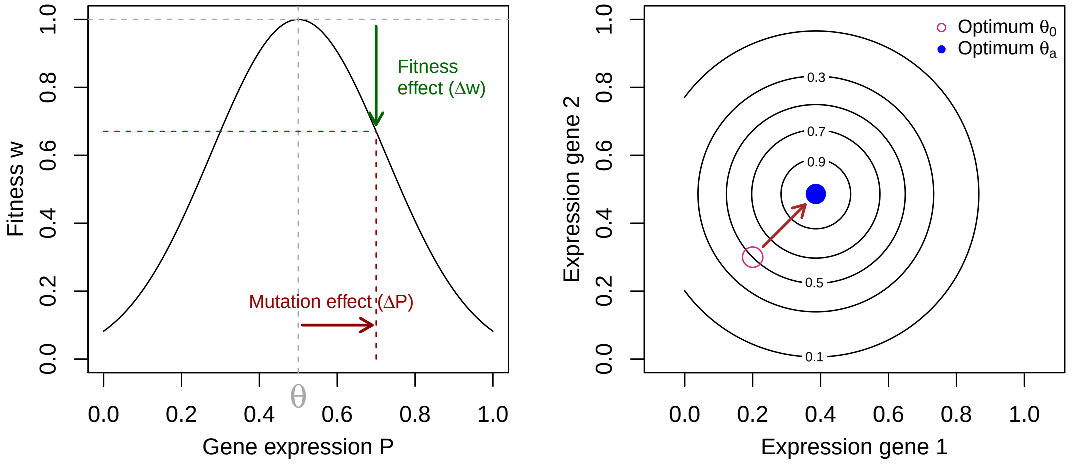


Supplementary Figure S1: **Graphical illustration of the fitness function**. **Left**: marginal fitness function (on a single gene). Selection is stabilizing around an optimum $\theta = 0.5$ in this case. The figure illustrates the effect of a mutation on an "optimal" genotype, that affects gene expression by a quantity $\Delta P$ (the mutational effect - on phenotype), and decreases the fitness by a quantity $\Delta w$ (the fitness effect of the mutation). In the simulations, 5 genes (out of 10 in the network) were submitted to this stabilizing selection. **Right**: To study the effect of mutations on the different topologies, 2 genes have their optima shifted by $\delta_\theta \simeq 0.186$, which decreases the fitness by 50%. Recovering the maximum fitness (brown arrow) thus implies to evolve the expression of 2 genes, leaving the expression of the 3 other selected genes unchanged. Contour lines highlight the new fitness function.

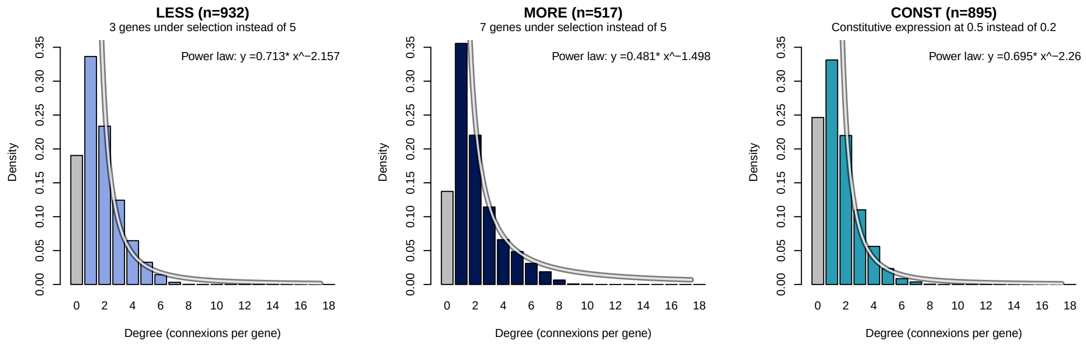


Supplementary Figure S2: **Different parameters also lead to the emergence of a scale-free topology**. We varied the radio of genes under selection 3 or 7 out of 10 genes (instead of 5 in the paper simulations), or the basal gene expression from 0.2 in the paper simulations to 0.5 here. The emergence of the scale-free topology appears conserved regardless.

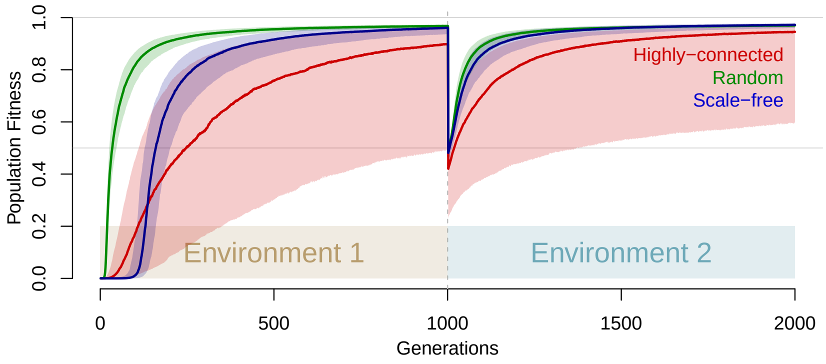


Supplementary Figure S3: **The initial conditions and topologies lead varying adaptation and re-adaptation dynamics**. This figure is equivalent to figure 2 in the main text, but includes both successful (population fitness > 0.95) and unsuccessful simulations.

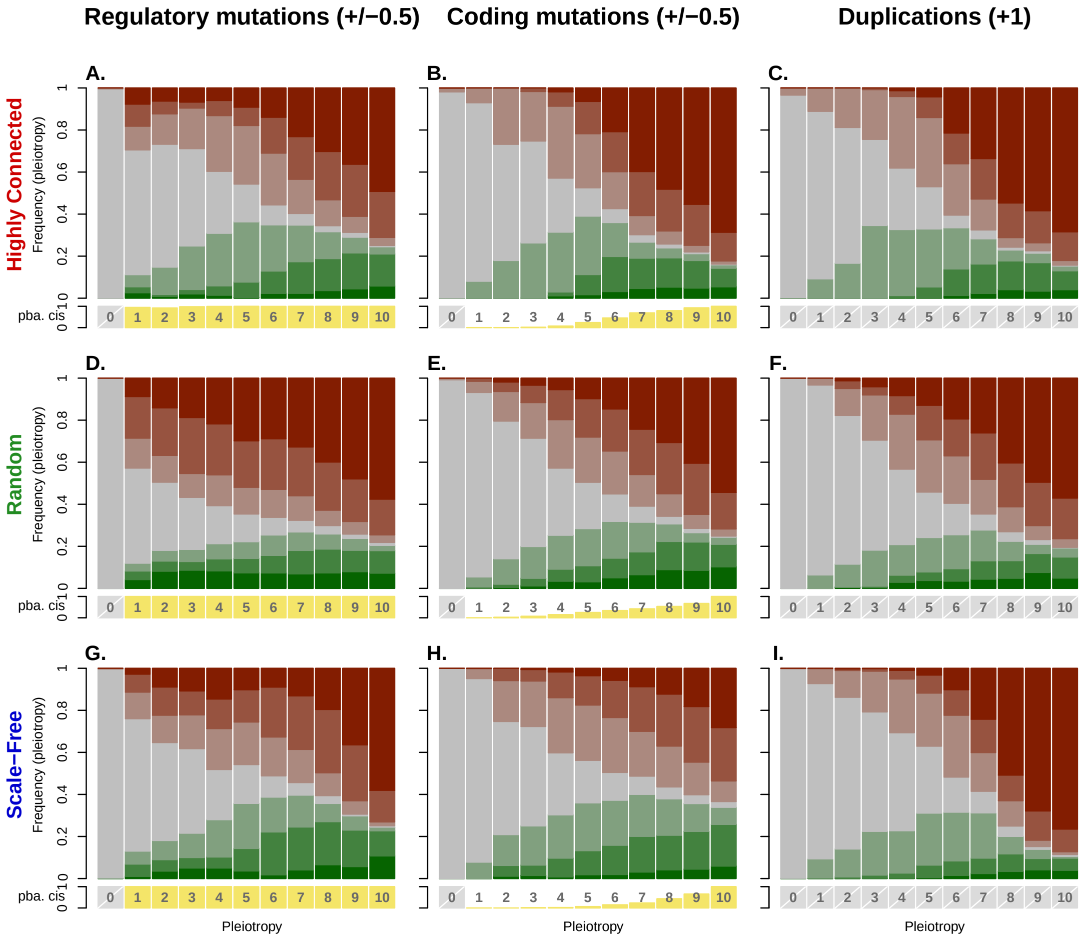


Supplementary Figure S4: **Alternative representation of Figure 4 (relative y axes).**

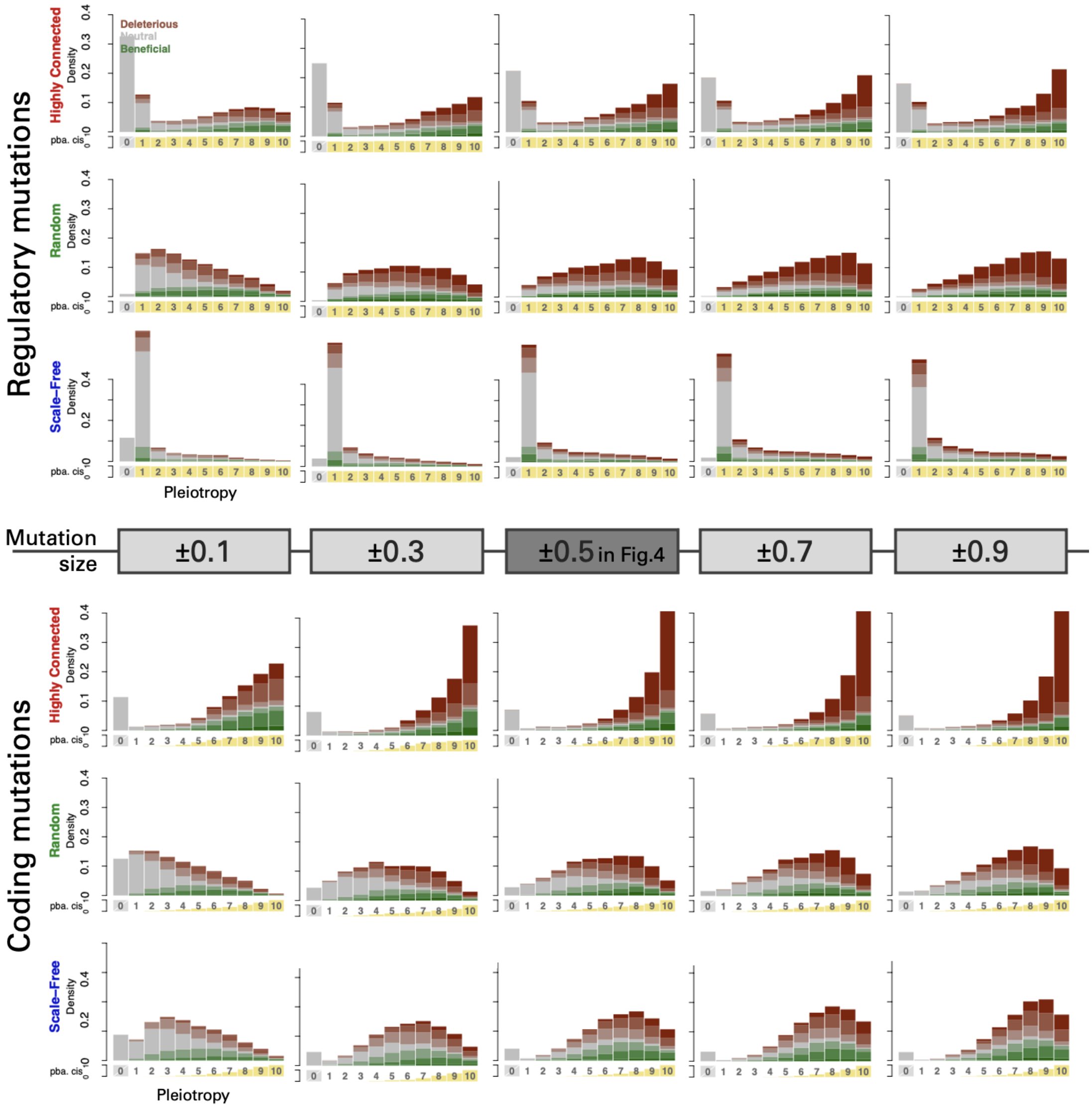


Supplementary Figure S5: **Pleiotropic and *cis*-acting effects for different mutation sizes** for regulatory and coding mutations. While a shift in pleiotropy is expected and observed, the overall distributions remain consistent across mutation sizes.

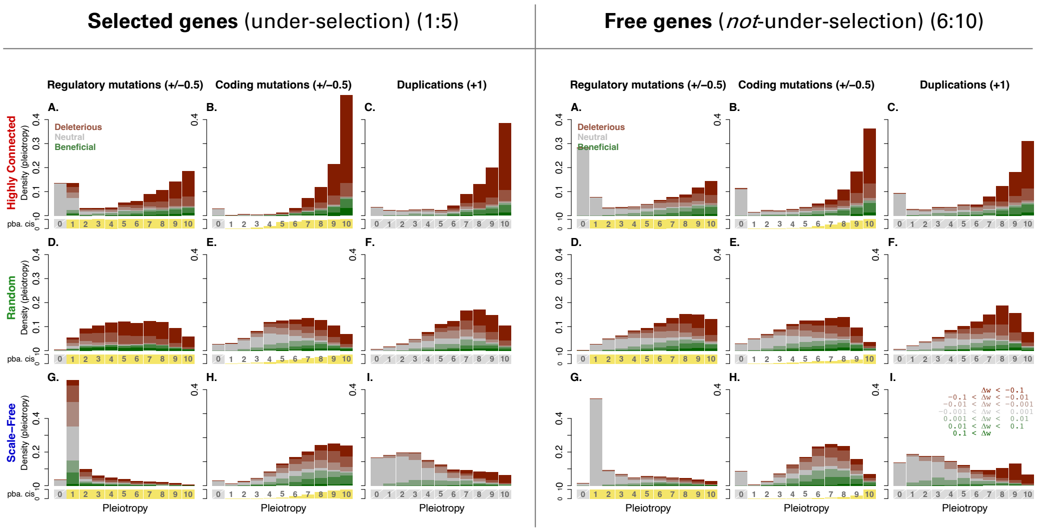


Supplementary Figure S6: **Pleiotropic and *cis*-acting effects for selected versus free genes**. We compared the pleiotropy of mutations affecting selected genes (**left**) with those affecting non-selected (free) genes (**right**) to analyze their distribution and fitness effects. Although the overall distributions are similar for both gene sets, the fitness effects differ, with free genes exhibiting a higher proportion of neutral mutations.

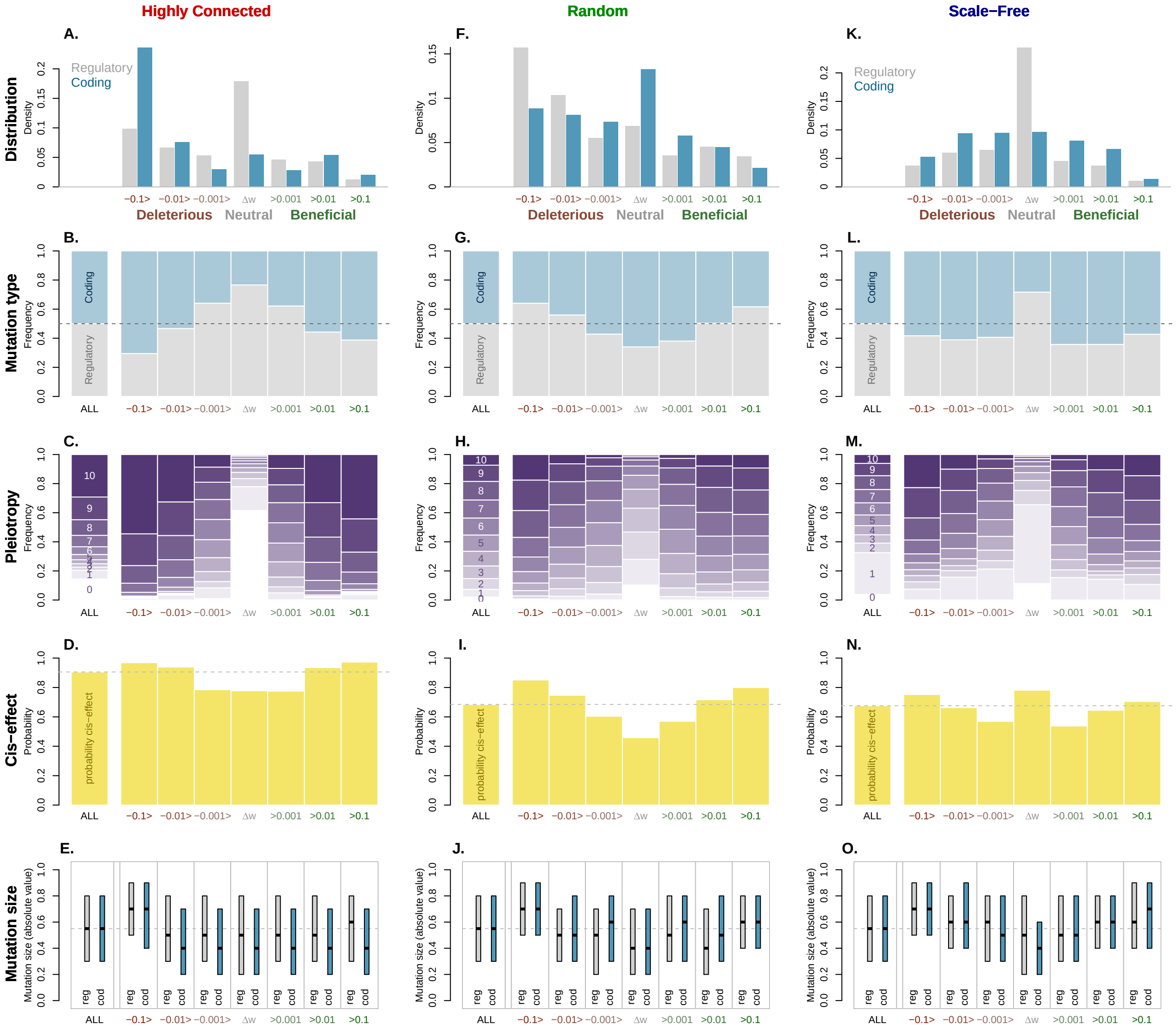


Supplementary Figure S7: **Enrichment in different types of mutations as a function of fitness effects**. This is the equivalent of figure 5, but for all three networks. (A, F, and K) distribution of fitness effects, (B, G, and L) coding vs. regulatory mutations, (C, H, M) pleiotropy, (D, I, N) *cis*- vs. *trans*-effects, (E, J, O) size of the mutation at the genetic level (boxes stand for median, q1 and q3).

### Supplementary movie S1

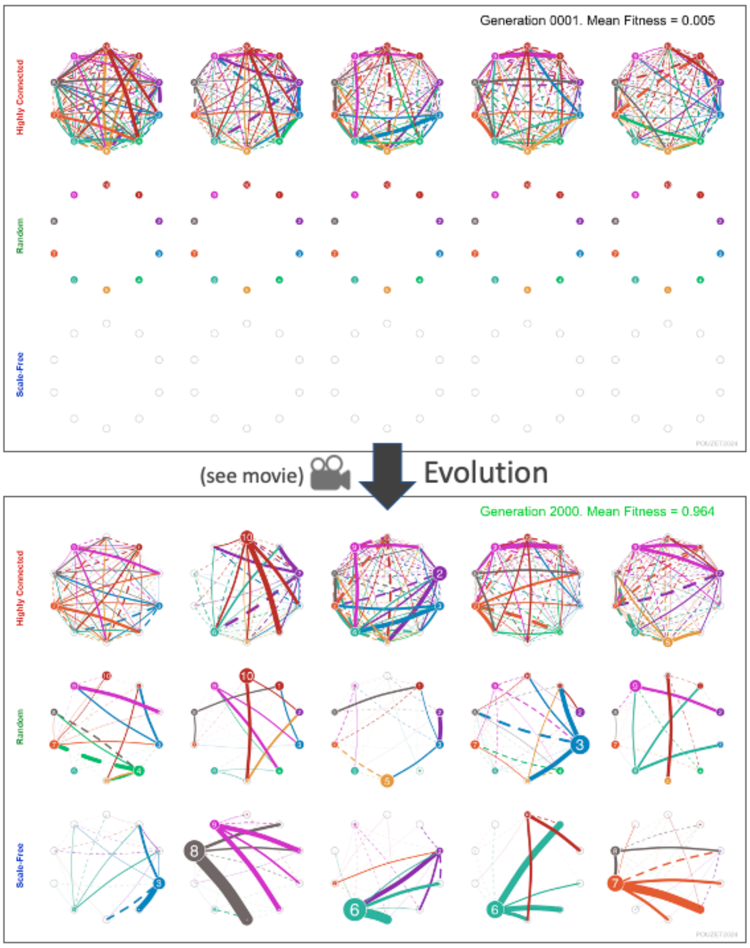


Evolution of networks from generation 1 to 2000, adapting to two successive environments, starting from three initial conditions and leading to three distinct topologies. After adaptation to the first environment (with genes 1 to 5 undergoing selection for a random expression optimum, while genes 6 to 10 evolved freely), an environmental shift occurs at generation 1000 (gene expression optima for two genes are altered), leading to a 0.5 fitness drop. Five networks are displayed for each of the three topologies examined in this study. The first (left) column includes networks used as representative examples in Figure 1. Each gene is color-coded, with edge colors indicating the regulatory gene: solid edges represent activation, and dashed edges represent repression. Node size reflects gene activity (”coding” value), and edge width represents the total strength of regulation. The movie is available on the article’s website or on the dedicated GitHub page: `https://github.com/spouze/GeneRation_Pouzet2024/`.

#### Supplementary movie S2

Zoomed-in, slower version of the movie S1 from generation 1 to 500. It shows the adaptation to the first environment, detailing the emergence of different topologies and their associated timings. These are the same images as in the previous movie S1, with the same corresponding legend.
